# A growth-based, high-throughput selection platform enables remodeling of 4-hydroxybenzoate hydroxylase active site

**DOI:** 10.1101/2020.05.11.088898

**Authors:** Sarah Maxel, Derek Aspacio, Edward King, Linyue Zhang, Ana Paula Acosta, Han Li

## Abstract

We report an aerobic, growth-based selection platform founded on NADP(H) redox balance restoration in *Escherichia coli*, and demonstrate its application in high-throughput evolution of oxygenase. A single round of selection enabled *Pseudomonas aeruginoasa* 4-hydroxybenzoate hydroxylase (PobA) to accept 3,4-dihydroxybenzoic acid efficiently, an essential step toward gallic acid biosynthesis. The best variant DA015 exhibited more than 5-fold higher catalytic efficiency compared to previously engineered enzymes. Structural modeling suggests precise re-organization of active site hydrogen bond network, which is difficult to obtain without deep navigation of combinatorial sequence space. We envision universal application of this selection platform in engineering NADPH-dependent oxidoreductases.

## Introduction

Oxygenases direct the strong oxidative power in molecular oxygen to activate inert C-H bonds with high regio- and stereo-specificity, which is difficult in traditional chemical synthesis^1^. This makes them valuable catalysts for production of industrially important and medicinal molecules such as ε-caprolactone^2^, omeprazole^3^, and artemisinin^4^. However, engineering oxygenases is challenging because of their highly complex catalytic mechanisms, which can obscure structure-function relationships^5^.

Recently, a high-throughput mass spectrometric enzyme activity assay was established and applied to P450 enzyme discovery^6^. However, throughput was still insufficient (~1000 variants screened for a protein library with a theoretical library size of 400) for libraries routinely generated in directed evolution (10^4^-10^6^ theoretical library size). To improve throughput, techniques using single cell or single enzyme analysis such as Fluorescence Activated Cell Sorting (FACS)^7^ and microfluidic assays^8^ have been developed. However, these methods are often limited by a need for colorimetric reactions^9^, substrate labeling^6^, or indirect activity readouts such as GFP fluorescence^10^. Additionally, these platforms often require extensive expertise and costly equipment.

We sought to address these limitations by developing a facile and universal selection scheme that employs *E. coli* growth as a simple readout. Specifically, we created an *E. coli* strain with overly high levels of NADPH/NADP^+^, which only grows on glucose in aerobic condition when the desired oxygenase activity is present to recycle NADPH. This selection scheme is inspired by Nature: previously, natural evolution was observed under high NADPH-stress which enabled *E. coli’s* respiratory chain to oxidize NADPH using oxygen^11,12^. This work is a significant departure from our past platform: the growth-based selection for engineering NADPH-dependent enzymes only functions in anaerobic conditions and hence is not compatible with oxygenases^13^.

We applied this selection platform for directed evolution of an NADPH-dependent monooxygenase *Pseudomonas aeruginosa* 4-hydroxybenzoate hydroxylase (PobA) to engineer high activity for a non-native substrate, 3,4-dihydroxybenzoic acid (3,4-DHBA)^14^, the precursor of a natural product antioxidant, gallic acid (GA). Notably, variants retained high activity with native substrate 4-hydroxybenzoate acid (4-HBA), which is also required in GA biosynthesis. Through a single round of selection of ~3×10^5^ variants from a four-residue, site-saturated mutagenesis library (theoretical library size 20^4^=1.6×10^5^), we obtained variants with roughly 5-fold improved catalytic efficiency (*k_cat_/K_M_*) for 3,4-DHBA compared to rationally designed variants reported previously^14^. Importantly, Rosetta modeling revealed that the selected PobA variants form intricate hydrogen bonding networks at the active site to recognize 3,4-DHBA and coordinate catalysis, which is difficult to recapitulate in engineered enzymes without extensive sampling of sequence space.

## Results and discussion

### Development and validation of the selection platform

The selection process relies on an engineered *E. coli* strain (MX203) with a growth deficiency linked to NADPH/NADP^+^ imbalance. The perturbed redox state results from deletion of central metabolism genes, glucose-6-phosphate isomerase *pgi* and phosphogluconate dehydratase *edd;* a critical rebalancing tool, soluble pyridine nucleotide transhydrogenase *udhA*; and a significant sink for reduced nicotinamide cofactors, NAD(P)H:quinone oxidoreductase *qor* (Figure 1A)^11^. These disruptions cause MX203 to exhibit poor growth in glucose minimal media (Figure S1) and we sought to restore growth through heterologous expression of NADPH-consuming enzymes. We employed water-producing oxidases, *Lb* NOX and *TP* NOX^15^ (on plasmids pLS101 and pLS102, respectively) (Figure 1B), derived from *Lactobacillus brevis.* When introduced into selection strain MX203, the NADPH-specific *TP* NOX restored growth, while the NADH-specific *Lb* NOX did not (Figure 1C). Desirable growth behavior demonstrated in both liquid and solid media provides flexibility in selection.

**Figure 1.**
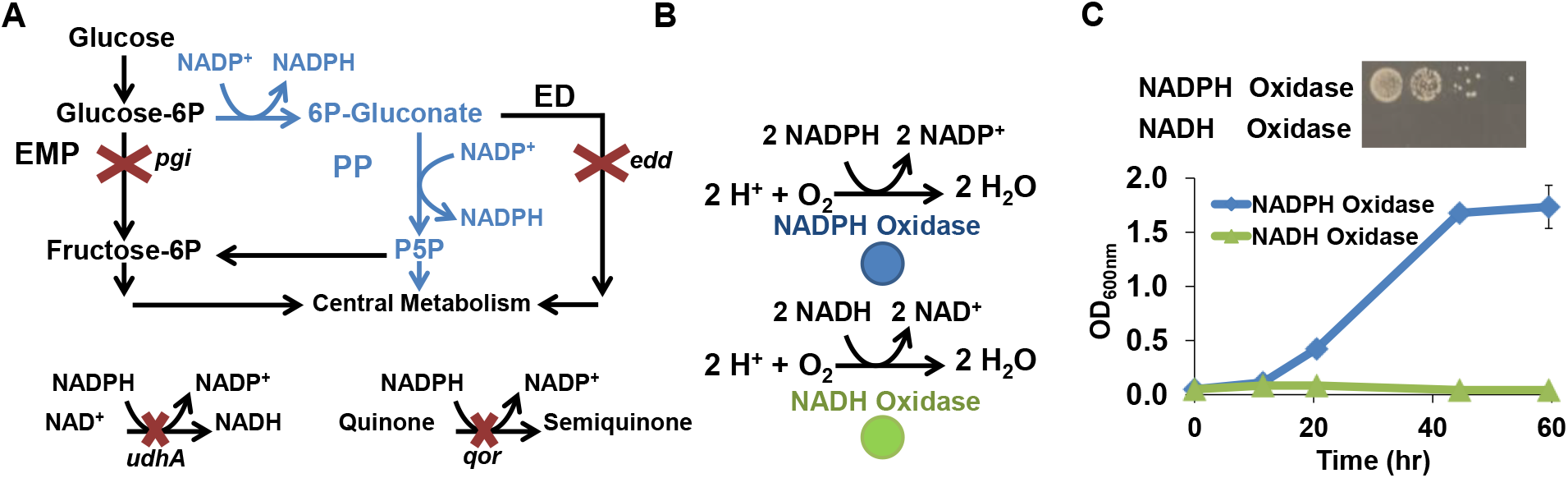
Development and validation of the selection strain with high NADPH accumulation. (A) Central metabolism re-direction through pentose phosphate (PP, blue) pathway and disruption of Embden-Meyerhof-Parnas (EMP) and Entner Doudoroff (ED) pathways increased NADP^+^ reduction. Deletion of NADPH re-oxidation transhydrogenase *udhA* and quinone oxidoreductase *qor* reduce NADPH sinks. (B) Complementary oxidases use reduced cofactors, NADPH (blue) or NADH (green), and oxygen to generate water. (C) In M9 minimal glucose media, growth restoration was achieved by heterologous expression of the NADPH-specific oxidase but not the NADH-specific oxidase in both solid and liquid media.

### Construction and selection of PobA library

Gallic acid has broad application in food and pharmaceutical industries^16^ and as the biosynthetic precursor of a platform chemical, pyrogallol^17^. Biosynthesis of gallic acid from simple carbon sources is hindered by the absence of a natural enzyme that can catalyze two consecutive hydroxylation steps, namely from 4-hydroxybenzoic acid (4-HBA) to 3,4-DHBA, and subsequently to gallic acid (Figure 2A). The NADPH-dependent 4-HBA hydroxylase from *P. aeruginosa* (*Pa* PobA) natively catalyzes the first step, and Chen and coworkers engineered additional activity for the second step^14^. While promising variants obtained from rational design, namely T385F and Y385F/T294A, enabled 3,4-DHBA hydroxylation, activity was not sufficient to prevent 3,4-DHBA accumulation during microbial production^17^. Furthermore, these mutations severely decreased the enzyme’s native activity required for the first hydroxylation step. With a high-throughput selection platform in hand, we sought to more systematically remodel the substrate binding pocket of *Pa* PobA.

**Figure 2.**
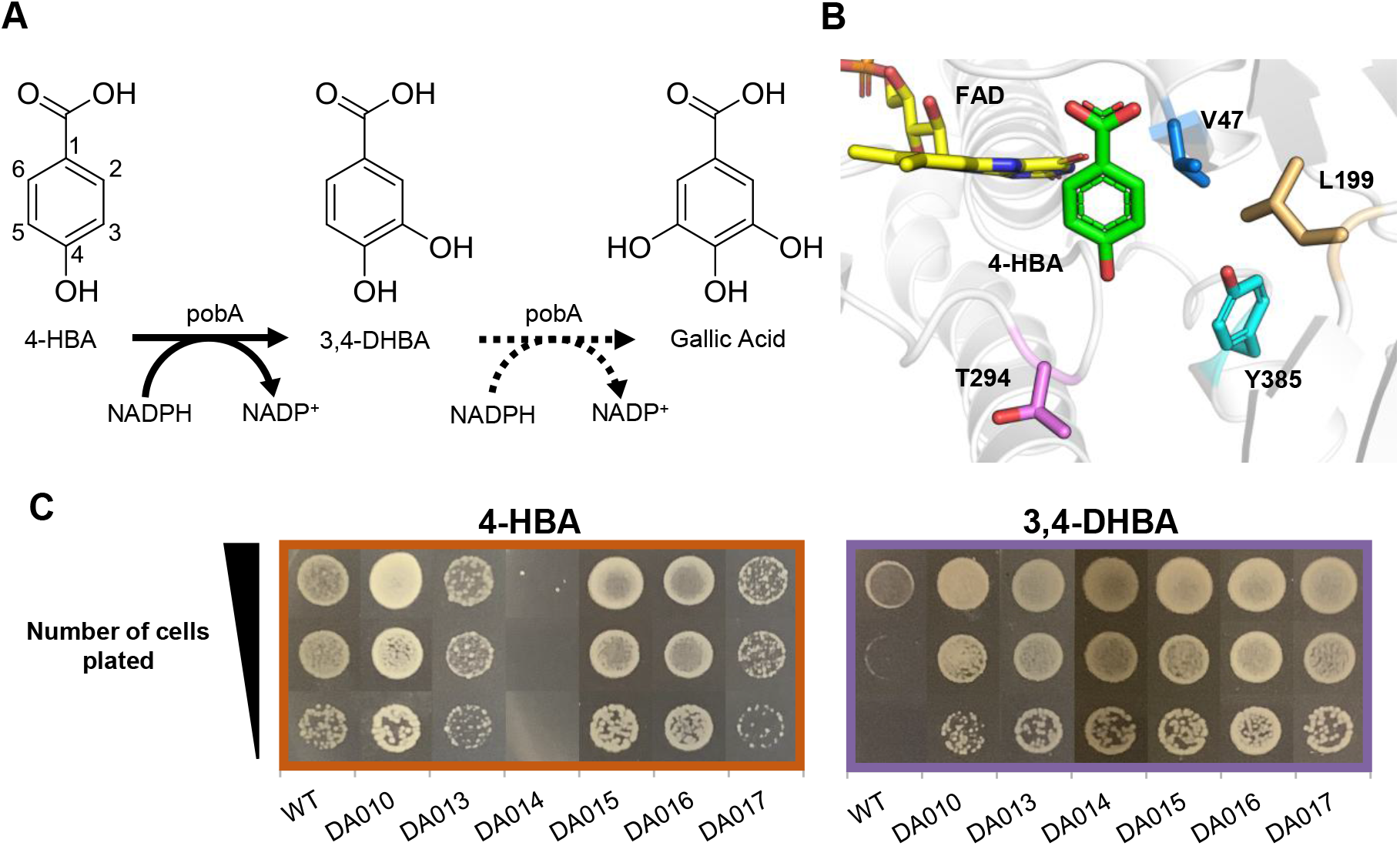
Construction and selection of the PobA library for 3,4-DHBA utilization. (A) Biosynthetic pathway of gallic acid involves two sequential hydroxylation steps. (B) Wild type *Pa* PobA crystal structure with 4-HBA bound (PDB: 1IUW). Residues highlighted were targeted for site-saturated mutagenesis. (C) Growth restoration phenotype for wild type PobA and variants identified in selection, with 4-HBA or 3,4-DHBA supplementation.

We hypothesized that achieving high activity with both 4-HBA and 3,4-DHBA would require the active site to flexibly accommodate the 3-hydroxyl of 3,4-DBA directly opposite the carbon 4a of flavin adenine dinucleotide (FAD) (Figure 2B). We inferred this mode of substrate binding would be more catalytically competent because oxygen transfer must occur between the flavin-C4a-hydroperoxide intermediate and a position on the aromatic ring with excess electron density, C5^18^. Guided by this hypothesis, we targeted three residues within 4 Å of the 3-hydroxyl for site-saturated mutagenesis (V47, L199, Y385 in 2B). Position T294 was selected for its proximity to the 4-hydroxyl of 4-HBA and potential role in the hydrogen bonding network of the active site^14^.

We simultaneously randomized residues 47, 199, 294, and 385 using NNK degenerate codons (Figure 2B). The resulting DNA library (pDA008) was transformed into selection strain MX203, yielding ~3×10^5^ independent transformants (see Methods in Supporting information), which was sufficient to cover 1.8-fold of the theoretical library size of 20^4^=1.6×10^5^. Selection was performed on agar plates with 2 g/L D-glucose in M9 minimal medium and 1 g/L 3,4-DHBA, at 30 °C for 60 hours. MX203 cells harboring wildtype *Pa* PobA served as a negative control. After incubation, ~600 colonies had formed on selection plates.

### Characterization of PobA variants obtained from selection

We picked eight colonies with robust growth and extracted the mutant *Pa* PobA plasmids. We verified growth restoration (Figure 2C) by re-transforming the plasmids into selection strain MX203 and characterizing growth with 3,4-DHBA or 4-HBA. The negative control, wild type PobA, showed little to no growth with 3,4-DHBA. Sequencing the eight candidates revealed only six unique residue combinations (Table S1), suggesting the high-throughput selection allowed comprehensive searching of the library to reach consensus. Discussion on observed consensus and trends in combinations is included in the supplementary information.

Four of the six unique variants were further characterized using purified proteins. All four variants gained significant improvements in 3,4-DHBA activity as measured *in vitro* by enzyme assay (Figure 3), indicating low frequency of false positives. Interestingly, DA014 which struggled to grow on plates containing 4-HBA, exhibited high specific activities for 4-HBA (Figure 3). As noted in previous work, this discrepancy might arise from differences between measured *in vitro* specific activity and *in vivo* activity^13^.

**Figure 3.**
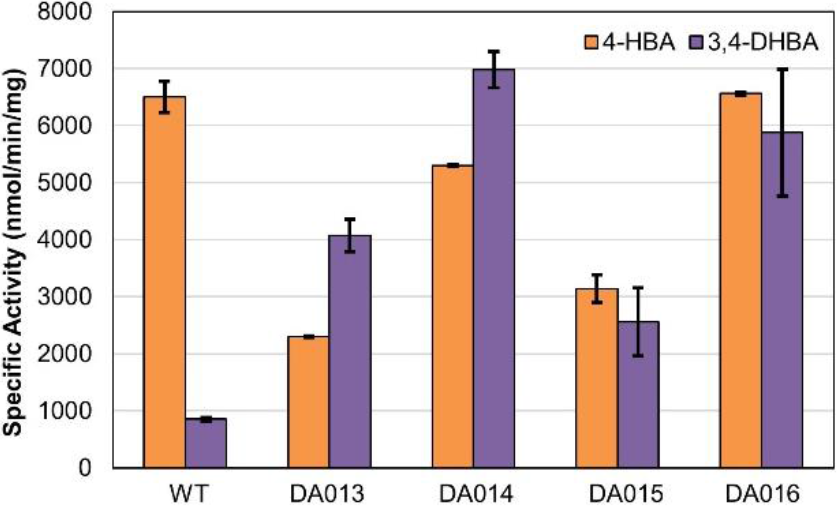
*In vitro* specific activity of four PobA variants identified in growth-based selection. All variants exhibited significantly improved activity towards 3,4-DHBA. Detailed assay conditions are described in the Methods section of Supporting information.

We further characterized two variants with fast growth rates with both 4-HBA and 3,4-DHBA, namely DA015 (L199R-T294C-Y385M) and DA016 (V47I-L199N-T294A-Y385I), alongside wild type *Pa* PobA (Figure 2C, Table 1). As shown in Table 1, catalytic efficiency (*k*_cat_/*K_M_*) of our best variant DA015, is 5.2-fold greater than previously engineered *Pa* PobA Y385F/T294A for 3,4-DHBA. To our knowledge, DA015 represents the most active 3,4-DHBA hydroxylase reported to date^14^. Importantly, DA015’s *K_M_* for 3,4-DHBA of ~48 μM, is 3-fold lower than that of PobA Y385F/T294A. Greatly improved affinity may be particularly beneficial for preventing 3,4-DHBA buildup during *in vivo* gallic acid biosynthesis by recognizing and rapidly converting 3,4-DHBA at low concentrations. Among all variants, DA016 has the highest *k_cat_* towards 3,4-DHBA (~4 s^-1^), which is over 50% of wild type PobA’s /e_cα t_ (~7.6 s^-1^) toward native substrate 4-HBA. This high turnover rate may be instrumental for *in vitro* catalysis. DA015 and DA016 also have 4-fold and 1.7-fold improved catalytic efficiency for 4-HBA compared to previously the reported PobA Y385F/T294A (Table 1). The 4-HBA hydroxylating activity of DA015 reached ~50% of that of the wild type enzyme.

**Table 1.**
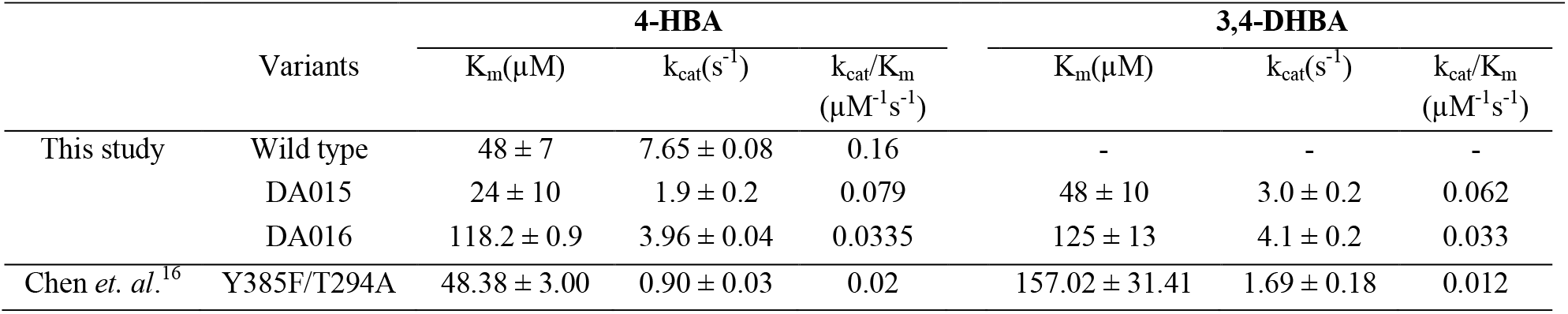
Kinetic characterization of PobA Variants

### Mechanistic study of the PobA variants by Rosetta modeling

We modeled the 3,4-DHBA binding pose with DA015 and DA016 to identify structural features contributing to enhanced activity over the wild type and previously reported variants. Wild type PobA crystal structure with 4-HBA bound (PDB: 1IUW) indicates 4-HBA binding mediated by a densely packed hydrogen bond network (Figure 4A). Previously reported PobA variants contained rationally designed mutations to disrupt hydrogen bond networks, aiming to increase backbone flexibility and substrate promiscuity^14,19^. In contrast, variants obtained in this study feature a remodeled hydrogen bond network tailored for 3,4-DHBA (Figure 4B, C).

**Figure 4.**
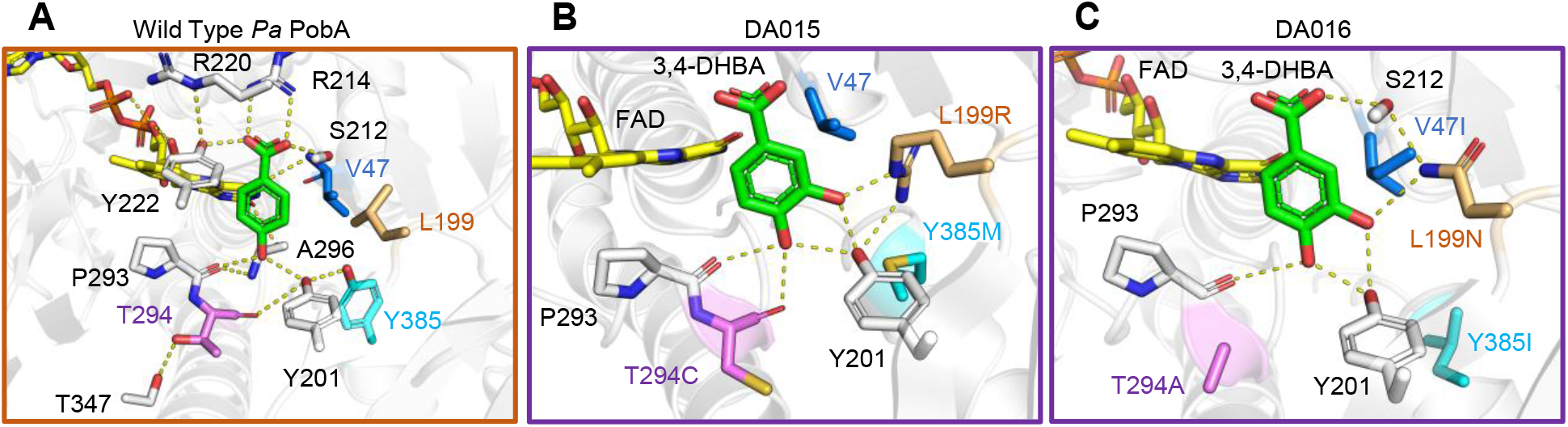
Models of 3,4-DHBA binding poses. (A) In wild type PobA, the hydrogen bonding networks stabilize 4-HBA and position the 3-carbon toward FAD for hydroxylation. (B) In DA015, L199R supports Y201 and forms a new contact to the ligand 3-hydroxyl. Y385M makes space, no substitution occurs at V47 which maintains close hydrophobic packing against L199R, and T294C loosens the helix for increased flexibility and improved backbone hydrogen bonding to the 4-hydroxyl. (C) In DA016, L199N forms interactions stabilizing S212 and to the ligand 3-hydroxyl. Y385I creates space, V47I braces L199N to minimize sidechain mobility, and T294A allows P293 to move closer to 3,4-DHBA. Importantly the mutations orient 3,4-DHBA such that the 5-carbon is optimally exposed to FAD for hydroxylation in both variants. Detailed methods of Rosetta modeling are included in the supporting information.

Specifically, in DA015 (L199R-T294C-Y385M), the L199R guanidinium extends to form two novel contacts, one to the Y201 hydroxyl and another to the 3,4-DHBA 3-hydroxyl; T294C eliminates a hydrogen bond to T347, allowing another new hydrogen bond to form between the backbone carbonyl to the 3,4-DHBA 4-hydroxyl (Figure 4B). In DA016 (V47I-L199N-T294A-Y385I), the L199N amide acts as a bridge stabilizing the ligand 3-hydroxyl and the S212 hydroxyl associated with the ligand carboxylic acid; V47I packs against and reinforces L199N; T294A, similarly to T294C, breaks a hydrogen bond to T347 and allows expanded loop mobility for improved backbone hydrogen bonding to the ligand 4-hydroxyl (Figure 4C). Comparison to wild type structure shows a binding pocket volume increase from 55 Å^3^ to 69 Å^3^ in DA015 and 72 Å^3^ in DA016 (Figure S2). The additional space accommodates the extra hydroxyl in 3,4-DHBA binding.

The discovered mutations restructure the central hydrogen bonding array for higher 3,4-DHBA binding affinity. Rational design to achieve the same effect is challenging due to the multiple, simultaneous substitutions required to reach this functionality. Designing hydrogen bonding networks is especially difficult due to their high cooperativity and sensitivity to small perturbations in length and angle between hydrogen bonding partners. For instance, singular L199 mutation to a polar residue contacting the ligand 3-hydroxyl would likely be ineffective without compensatory substitutions at V47 and Y385 relieving steric clash and reinforcing optimal hydrophobic packing. This highlights the effectiveness of our high-throughput selection approach in comprehensively exploring the sequence space and discovering highly synergistic mutations.

## Conclusion

In summary, we established a high throughput screening platform with potential applications for directed evolution of various NADPH-dependent oxygenases. This aerobic *in vivo* system demonstrated strict cofactor dependence and simple growth-based selection. We successfully engineered PobA for enhanced activity towards the non-native substrate 3,4-DHBA. In one round of selection, we identified a variant, DA015, with greatly improved activity for both 4-HBA and 3,4-DHBA compared to variants previously designed with restricted exploration of the available sequence space. Rosetta modeling suggests enhanced activity is attributed to concerted changes in binding pocket volume, shape complementarity, and more importantly, formation of a fully connected hydrogen bond network in the active site. Because NADPH provides reducing power for a variety of oxygen-dependent industrially important enzyme classes such as Bayer-Villiger monooxygenases and cytochrome P450s, we envision applications of this platform for engineering a broad range of catalysts.

## SUPPORTING INFORMATION

Experimental methods, plasmids and strains used in this study (Table S1), detailed analysis of the sequences of PobA variants obtained from selection (Table S2), Growth phenotypes of *E. coli* strains with modified NADPH metabolism (Figure S1), and active site volumes of wild type, DA015, and DA016 PobA (Figure S2).

## AUTHOR CONTRIBUTION

S.M., D.A., and H.L. designed the experiments, S.M., D.A., L.Z., and A.P.P performed the experiments and analyzed the results, E.K. performed Rosetta modeling, All authors wrote the manuscript.

## NOTES

The authors declare no competing financial interest.

## ACKNOWLEDGMENT

H. L. acknowledges support from University of California, Irvine, the National Science Foundation (NSF) (award no. 1847705), and the National Institutes of Health (NIH) (award no. DP2 GM137427). S.M. acknowledges support from the NSF Graduate Research Fellowship Program (grant no. DGE-1839285). D.A. acknowledges support from the Federal Work Study Program funded by the U.S. Department of Education.

