## Supporting information for "A growth-based, high-throughput selection platform enables remodeling of 4-hydroxybenzoate hydroxylase active site"

**A. Methods**

**B. Supplemental Table and Figures**

Table S1 Strains and Plasmids used in this study

Table S2 Sequences of PobA variants obtained in the selection

Figure S1 Modified metabolisms of *E. coli* strains displayed distinct growth phenotypes in M9 Seletion Media with glucose as carbon source

Figure S2 PobA variants show greater active site volumes compared to the wild type, and are capable of accepting the bulkier 3,4-DHBA

**C. References**

### A. Methods

**Media and Growth Conditions**. Cloning was carried out with *E. coli* XL-1 blue and protein expression was performed with *E. coli* BL21 (DE3). All *E. coli* were cultured in 2xYT containing 16 g/L Tryptone, 10 g/L Yeast Extract and 5 g/L NaCl unless otherwise noted. M9 Wash Buffer consisted of 1 mM $MgSO_4$, 0.1 mM $CaCl_2$, trace metal mix A5 with Co ($H_3BO_3$ 2860 μg/L $H_3BO_3$ 2860 μg/L, $MnCl_2 \cdot 4H_2O$ 1810 μg/L, $ZnSO_4$ $7H_2O$ 222 μg/L, $Na_2MoO_4$, $2H_2O$ 390 μg/L, $CuSO_4$, $5H_2O$ 79 μg/L, $Co(NO_3)_2$ $6H_2O$ (49 μg/L), and BD Difco M9 salts ($Na_2HPO_4$ 6.78 g/L, $KH_2PO_4$ 3g/L, NaCl 0.5 g/L, $NH_4Cl$ 1 g/L). M9 Selection Media shared the same composition of M9 Wash Buffer with the inclusion of 2 g/L D-glucose, 0.01 g/L thiamine, 0.04 g/L $FeSO_4 \cdot 7H_2O$. For solid media M9 Selection Plates, 15 g/L agar was added in addition to M9 Selection Media composition. Substrates 3,4-DHBA or 4-HBA were added at a concentration of 1 g/L prepared from a DMSO stock when appropriate. Concentrations for antibiotic selection were 100 mg/L for ampicillin, 50 mg/L for kanamycin, and 50 mg/L for spectinomycin. All strains were cultured at 37 ℃ with 250 rpm agitation. Induction was initiated with final concentrations of 0.1% arabinose for strains with $P_{BAD}$ promoter, and 0.1 mM IPTG for strain with $P_{lac}$ promoter.

**Strain Construction.** *E. coli* strain JW3985-1 (*Δpgi::kan*) served as starting strain for MX203 construction. Plasmid pCP20 was used to eliminate kanamycin resistance in between knockouts performed[1]. Knockouts Δ*qor* and Δ*edd* on Δ*pgi* were generated using the P1 phage transduction method[2]. Keio collection strains[1] JW4011-2 (*Δqor::kan*) and JW1840-1 (*Δedd::kan*) served as donors for the generation of P1 lysate with gene knockout cassettes containing a kanamycin resistance marker. Knockout of *udhA* on *ΔpgiΔeddΔqor::kan* was performed with λ red recombinase system using pKD46[1]. Nested PCR was used to generate kanamycin cassette with 80 bp of homologous DNA on each side of *udhA* recombination site. All knockouts were confirmed by MyTaq Colony PCR (Bioline).

**Plasmid Construction.** All PCR fragments were generated using PrimeSTAR Max DNA Polymerase (TaKaRa) unless otherwise noted. The *Lactobacillus brevis nox* gene was amplified from genomic DNA from *Lactobacillus brevis* strain 118-8 [ATCC® 367™]. After PCR and gel extraction, the *Lb nox* gene fragment was inserted into the vector backbone (pRSF ori, $Spec^r$) using Gibson isothermal DNA assembly method[3], resulting in plasmid pLS101. Plasmid pLS102 carrying the *TP nox* gene was generated using plasmid pLS101 as a template and using site directed mutagenesis to introduce the following mutations G159A-D177A-A178R-M179S-P184R.

The *Pseudomonas aeruginosa pobA* gene was amplified from *Pseudomonas aeruginosa* PAO1-LAC [ATCC® 47085™] genomic DNA. After PCR and gel extraction, the *pobA* gene fragment was inserted into pQElac vector backbone which contains a 6×His tag at the N-terminus (ColE1 ori, $Amp^r$) using Gibson isothermal DNA assembly method, resulting in plasmid pDA006.

**PobA Library Construction.** All PCR reactions with degenerate primers were generated with KOD Xtreme Hot Start DNA Polymerase (Novagen) as well as through splicing by overlap extension (SOE) PCR. To construct the *Pa pobA* NNK library a forward primer containing an NNK codon at position V47 was used in a PCR reaction with a reverse primer containing an MNN codon at position L199 to generate the first fragment. A forward primer that begins at the codon for position I200 was used in a PCR reaction with a reverse primer that begins at the codon for position P293 to generate the second fragment. A third fragment was generated by PCR reaction of a forward primer containing an NNK codon at position T294 with a reverse primer containing an MNN codon at position Y385. All three fragments were gel purified and used as templates in a SOE PCR reaction to generate an insert that contained degenerate codons at the four sites of interest. The gel purified insert fragment and plasmid pDA006 were separately digested with EagI-HF and XbaI. Both digestion products were again gel purified and the appropriate length fragments were assembled by ligation, purified, and transformed into ElectroMAX DH10β cells (Invitrogen) using electroporation. Then cells were rescued with SOC medium at 37 °C for 1 hour and then added to 20 ml 2xYT medium with appropriate antibiotics. 2 μL, 20 μL, and 200 μL of culture was sampled from the culture and plated on an 2xYT agar plates with appropriate antibiotics. The remaining liquid culture was incubated at 37 °C with 250 r.p.m. agitation for 10 hours, before extraction of the library DNA. The plates were incubated at 37 °C overnight, and colonies formed were counted to estimate the library size. 10 single colonies were also cultured individually to extract plasmids, which were sequenced to sample the sequence space of the library. All 10 plasmids sequenced contained unique mutation patterns in the four targeted sites and no other unintended mutations were present.

**Growth Rescue Conditions.** The growth rescue condition used for Figure 1C was as follows: Briefly, the strains tested were first cultured in 2xYT under aerobic conditions at 30 °C overnight with appropriate antibiotics and inducers. Next, overnight cultures were washed 3 times and re-suspended in M9 Wash Buffer. For solid growth, serial dilutions of $10^6$ cells/mL, $10^5$ cells/mL, $10^4$ cells/mL, and $10^3$ cells/mL were prepared in M9 Wash and 2 μL aliquots were dispensed in series on an agar plate of M9 Selection Media, with appropriate antibiotics and inducers. Plates were grown at 30 °C and photos were taken to document growth progress. For liquid growth, a 0.1% (v/v) volume of washed culture was used to inoculate to an $OD_{600nm}$ of ~0.02 in 0.3 mL of M9 Selection Media, with appropriate antibiotics and inducers. Culture tubes were incubated at 30 °C in a rotary shaker, and OD600 nm was measured using a 96-well plate reader.

**Transformation of PobA library by Electroporation of MX203.** To generate electro-competent cells of *E. coli* MX203, cells were cultured in 200 mL SOB medium with appropriate antibiotics at 30 °C with shaking at 250 r.p.m. until $OD_{600nm}$ reached 0.4-0.6. The culture was chilled on ice for 15 min and the cells were pelleted at 4 °C, 4000 ×g. The cells were washed at 4 °C three times with 40 mL 10% glycerol in water (sterile, ice cold). After, cells were finally resuspended with 500 μL 10% glycerol in water (sterile, ice cold), and aliquoted for transformation.

The transformation was performed as follows: After MX203 electro-competent cells were thawed on ice, 20 uL of library pDA008 was added to 200 μL competent cells. Cell-DNA mixture was added to four ice chilled 1 mm gap electroporation cuvettes (55 μL of per electroporation cuvette). Cells were electroporated at 2 kV, 129 Ω, 50 μF, resistance 2.5 kV; 200 uL of SOC medium immediately added and transferred to a microcentrifuge tube at room temperature. This step was repeated twice more. Cells were rescued at 37 °C with shaking for 1 hour. Serial dilution of the cells was performed and then plated on 2xYT agar plates with appropriate antibiotics. After incubation at 37 °C overnight, colonies formed were counted to estimate transformation efficiency.

**Selection of *Pa* PobA Library.** *E. coli* MX203 was transformed with the *Pa* PobA library pDA008 by electroporation. After rescue in SOC medium for 1 hour, cells containing plasmid pDA008 were combined in 20 mL 2xYT with appropriate antibiotics in a 250 mL baffled shake flask. Controls were added to 5 mL 2xYT in a 50 mL conical tube (cap loose to allow increased aeration) and grown at 30 °C for ~7-8 hours or until $OD_{600nm}$ = 0.6 was reached. Subsequently, controls and the library were induced by addition of IPTG. Cultures were grown for an additional 4 hours or until $OD_{600nm}$ = ~1.68.

To prepare cells for the selection condition, 1 mL of each culture was pelleted in 2 mL microcentrifuge tubes and washed three times with M9 Wash Buffer.

After wash, cells were finally re-suspended in 1 mL M9 Wash Buffer. Cells were diluted with M9 Wash Buffer to a final cell concentration of ~$10^7$ cells/mL. 100 µL of this cell suspension was plated on M9 Selection Plates with supplemented with either 3,4-DHBA or 4-HBA and incubated at 30 ℃ for 60 hours. The colonies were monitored every 12 hours. The colonies were picked and re-streaked onto the same type of plates and again incubated at 30 ℃ to obtain single colonies. Single colonies were cultured in liquid media to extract plasmids using QIAprep Spin Miniprep kit (Qiagen) to yield pDA010, pDA011, pDA012, pDA013, pDA014, pDA015, pDA016, and pDA017.

**Protein Expression and Purification of *Pa* PobA Variants.** Plasmids pDA006, pDA013, pDA014, pDA015, pDA016 *E. coli* were transformed into BL21 (DE3) and cultured in 2xYT medium with appropriate antibiotics at 37 ℃ overnight. Overnight cultures were then used to inoculate 20 mL 2xYT medium with antibiotics at 37 ℃, agitated at 250 r.p.m. to an $OD_{600nm}$ = 0.5. The cultures were induced with 0.5 mM IPTG and incubated at 30 ℃ (250 r.p.m) for 24 h. Recombinant protein was purified using His-Spin Miniprep kit (Zymo Research Corporation) according to the manufacturer's instructions. The concentrations of purified proteins were quantified by Bradford assay against a BSA standard curve.

**Characterization of *Pa* PobA Variants.** Protein was purified as detailed above. The enzyme activity was measured using previously reported methods[4]. Specific activity measurements were started by the addition of purified protein to the assay mixture (200 µl) containing 100 mM Tris-HCl (pH 8.0), 10 µM FAD, 1 mM NADPH, 1 mM substrate (4-HBA or 3,4-DHBA). For kinetic parameters measurement, reactions were started by the addition of NADPH to the assay mixture (100 µL) after 10 min preincubation at 30 ℃ containing 100 mM Tris-HCl (pH 8.0), 10 µM FAD, 1 mM NADPH, and 0-1000 µM substrate. For wild type PobA, DA015, and DA016, the enzyme concentrations were 60 nM, 120 nM, and 120 nM respectively. The assay was conducted at 30 ℃ and slopes were calculated from the first 2 min of reaction and absorbance at 340 nm was measured in 10 s intervals. Kinetic parameters were estimated through non-linear regression of the Michaelis-Menten equation. Activity assays were conducted in duplicate.

**Rosetta modeling.** Docking models for DA015 and DA016 with bound 3,4-DHBA and FAD were generated with Rosetta[5]. The crystal structure of PobA with FAD and 4-HBA bound (PDB: 1IUW) was used as the starting template[6]. The structure model for 3,4-DHBA was downloaded from the PubChem database, and the starting pose was selected by alignment onto 4-HBA[7]. The Rosetta docking protocol involved mutation from the wild type structure, repeated rounds of random rigid-body perturbation by translation and rotation for 3,4-DHBA, and optimization of active site rotamers through sidechain repacking and minimization. All moves were sampled with the Monte Carlo method, no coordinate restraints were utilized, and full flexibility for ligand and protein backbone torsions was allowed to identify the optimal binding pose. A total of 1,000 docking trials was run for each variant, the top 100 models based on total Rosetta energy were sorted on interface energy scores and the model with the most favorable interface energy to 3,4-DHBA was selected as the reference. Analysis of binding pocket volume was completed with POVME 3.0 and protein figures were generated with PyMol [8] (Schrödinger LLC, 2020).

**B. Supplemental Table and Figures**

**Table S1. Strains and Plasmids used in this study**

| **Strains** | **Description** | **Reference** |
|---|---|---|
| XL-1 blue | Cloning strain | Stratagene |
| BL21 (DE3) | Protein expression strain | Invitrogen |
| BW25113 | *E. coli* F-, DE(araD-araB)567, lacZ4787(del)::rrnB-3, LAM-, rph-1, DE(rhaD-rhaB)568, hsdR514 | Datsenko *et al.*[1] |
| DH10β | Electrotransformation strain | Invitrogen |
| JW3985-1 | BW25113 Δ*pgi::kan* | Coli Genetic Stock Center |
| JW4011-2 | BW25113 Δ*qor::kan* | Coli Genetic Stock Center |
| JW1840-1 | BW25113 Δ*edd::kan* | Coli Genetic Stock Center |
| MX201 | BW25113 Δ*pgi* Δ*edd* | This study |
| MX202 | BW25113 Δ*pgi* Δ*edd* Δ*qor* | This study |
| MX203 | BW25113 Δ*pgi* Δ*edd* Δ*qor* Δ*udhA::kan* | This study |

| **Plasmids** | **Description** | **Reference** |
|---|---|---|
| pCP20 | Temperature-inducible yeast Flp recombinase gene controlled by λ*c*I*ts857* in a temperature-sensitive replicon | Datsenko *et al.*[1] |
| pKD46 | *ori101 repA101ts* $P_{BAD}$ *gam-bet-exo* $Amp^r$ | Datsenko *et al.*[1] |
| pQElac | $Amp^r$ ; ColE1 ori; $P_{LlacO1}$. Expression vector | Li *et al.*[9] |
| pLS101 | pRSF $P_{BAD}$::*Lb nox*, $Spec^r$ | This study |
| pLS102 | pRSF $P_{BAD}$::*Tp nox* (*Lb nox* G159A-D177A-A178R-M179S-P184R), $Spec^r$ | This study |
| pDA006 | pQElac 6xHis *Pa pobA*, $Amp^r$ | This study |
| pDA008 | pQElac 6xHis *Pa pobA* V47-L199-T294-Y385 NNK library, $Amp^r$ | This study |
| pDA010 | pQElac 6xHis *Pa pobA* L199V-T294C-Y385I, $Amp^r$ | This study |
| pDA011 | pQElac 6xHis *Pa pobA* V47L-L199N-T294A-Y385L, $Amp^r$ | This study |
| pDA012 | pQElac 6xHis *Pa pobA* -L199R-T294C-Y385M, $Amp^r$ | This study |
| pDA013 | pQElac 6xHis *Pa pobA* V47V-L199I-T294C-Y385W, $Amp^r$ | This study |
| pDA014 | pQElac 6xHis *Pa pobA* V47L-L199N-T294A-Y385L, $Amp^r$ | This study |
| pDA015 | pQElac 6xHis *Pa pobA* L199R-T294C-Y385M, $Amp^r$ | This study |
| pDA016 | pQElac 6xHis *Pa pobA* V47I-L199N-T294A-Y385I, $Amp^r$ | This study |
| pDA017 | pQElac 6xHis *Pa pobA* L199I-T294V-Y385W, $Amp^r$ | This study |

Abbreviations indicate source of genes: *Lb*, *Lactobacillus brevis*; *Pa*, *Pseudomonas aeruginosa*

**Table S2. Sequences of PobA variants obtained from the selection**

| | Val 47 | | Leu 199 | | Thr 294 | | Tyr 385 | |
|---|---|---|---|---|---|---|---|---|
| | Codon | Residue | Codon | Residue | Codon | Residue | Codon | Residue |
| WT | GTG | V | CTG | L | ACC | T | TAT | Y |
| DA010 | GTG | V | GTG | V | TGT | C | ATT | I |
| DA011 | TTG | L | AAT | N | GCG | A | TTG | L |
| DA012 | GTT | V | CGT | R | TGT | C | ATG | M |
| DA013 | GTG | V | ATT | I | TGT | C | TGG | W |
| DA014 | TTG | L | AAT | N | GCG | A | CTG | L |
| DA015 | GTT | V | CGT | R | TGT | C | ATG | M |
| DA016 | ATT | I | AAT | N | GCT | A | ATT | I |
| DA017 | GTT | V | ATT | I | GTG | V | TGG | W |

Sequencing of the eight candidates revealed six unique residue combinations at the mutation sites. Based on the sequencing results we observed consensus at individual sites or weak trends in their combinations. For example, as shown in Table S1, all eight mutants have a hydrophobic branched-chain amino acid at position V47 (valine, leucine, or isoleucine), and all eight residues observed at position T294 have side chains that do not participate in hydrogen bonding as predicted for hydrogen bond network flexibility by Chen *et al*[4]. Positions L199 and Y385 follow a loose trend of size complementarity and hydrogen bond character. Mutations at these positions appear to converge to combinations with a single hydrogen bond acceptor with potentially cooperative mutations in the adjacent positions to accommodate unexpectedly large residues such as tryptophan and arginine.

Interestingly, two variants, DA011 and DA014, struggled to grow on the enzyme's native substrate, 4-HBA. Sequencing revealed that these two variants encode the same mutations (V47L, L199N, T294A, Y385L), but with different codon usage at position Y385L which suggests consensus from a diverse sequence library, as shown in Table S1.

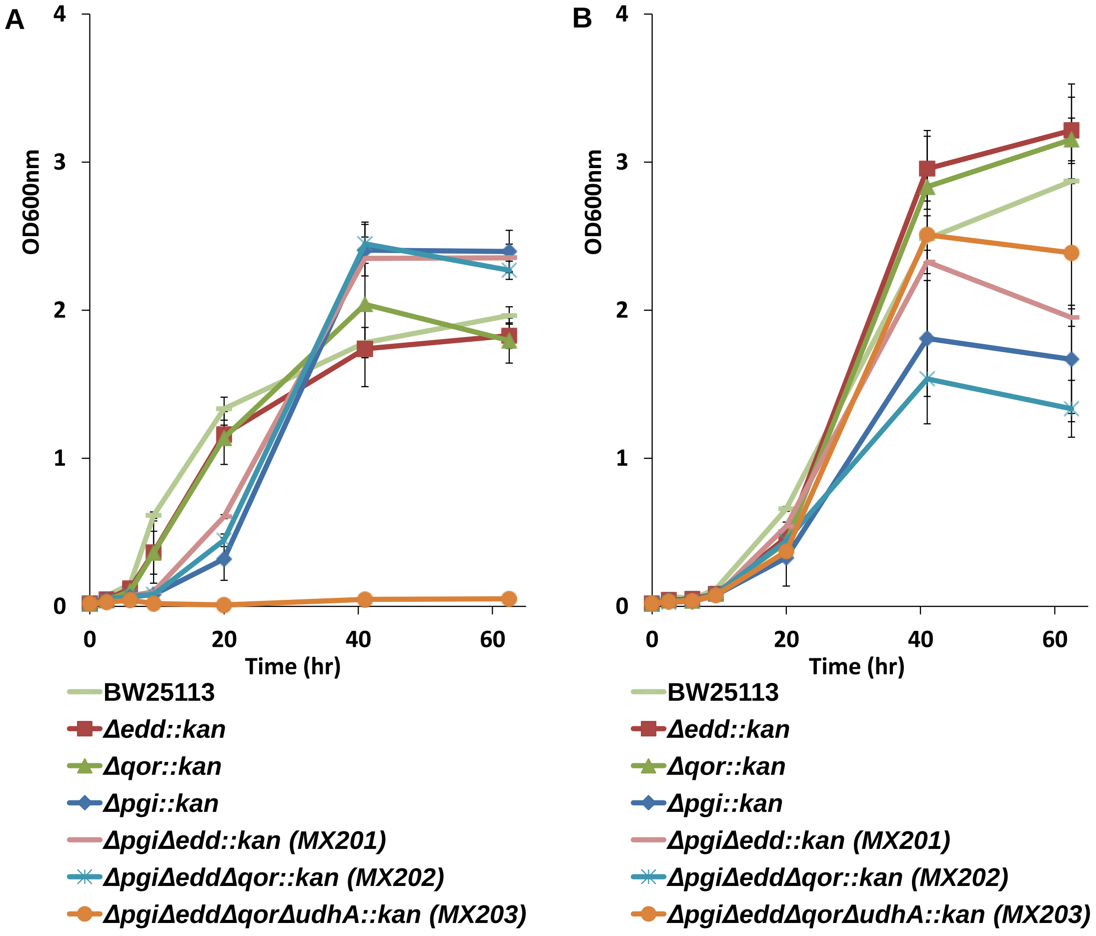


**Figure S1. Modified metabolisms of *E. coli* strains displayed distinct growth phenotypes in M9 Selection Media with glucose as carbon source.** (A) In M9 minimal media with glucose as the sole substrate growth defects observed in modified strains with deletion of *pgi*. Deletion of *udhA* in strain MX203 drastically enhanced growth defects compared to MX202 which did not have the *udhA* deletion. (B) With glycerol as the sole substrate. Growth defects observed were minimal, however a wide range of final culture densities was observed.

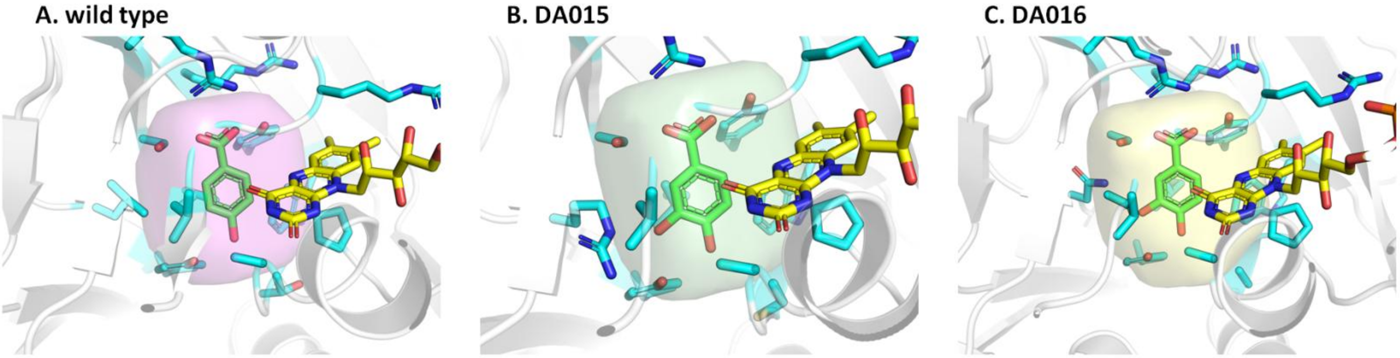


**Figure S2. PobA variants show greater active site volumes compared to the wild type, and are capable of accepting the bulkier 3,4-DHBA.** Binding pocket volumes calculated with POVME. A) wild type PobA has binding pocket volume 55 Å$^3$ B) DA015 has binding pocket volume 69 Å$^3$ C) DA016 has binding pocket volume 72 Å$^3$
